# Emergence and evolution of antimicrobial resistance genes and mutations in *Neisseria gonorrhoeae*

**DOI:** 10.1101/2020.10.26.354993

**Authors:** Koji Yahara, Kevin C. Ma, Tatum D. Mortimer, Ken Shimuta, Shu-ichi Nakayama, Aki Hirabayashi, Masato Suzuki, Michio Jinnai, Hitomi Ohya, Toshiro Kuroki, Yuko Watanabe, Mitsuru Yasuda, Takashi Deguchi, Vegard Eldholm, Odile B. Harrison, Martin C. J. Maiden, Yonatan H. Grad, Makoto Ohnishi

## Abstract

Antimicrobial resistance in *Neisseria gonorrhoeae* is a global health concern. Strains from two internationally circulating sequence types, ST-7363 and ST-1901, have acquired resistance to treatment with third-generation cephalosporins mainly due to the emergence of mosaic *penA* alleles. These two STs were first detected in Japan; however, when and how the mosaic *penA* alleles emerged and spread to other countries remains unknown. Here, we addressed the evolution of *penA* alleles by obtaining complete genomes from three Japanese ST-1901 clinical isolates harboring mosaic *penA* allele 34 (*penA-*34) dating from 2005 and generating a phylogenetic representation of 1,075 strains sampled from 37 countries. We also sequenced the genomes of 103 Japanese ST-7363 *N. gonorrhoeae* isolates from 1996-2005 and reconstructed a phylogeny including 88 previously sequenced genomes. Based on an estimate of the time of emergence of ST-1901 harboring mosaic *penA-34* and ST-7363 harboring mosaic *penA*-10, and >300 additional genome sequences of Japanese strains representing multiple STs isolated in 1996-2015, we suggest that *penA*-34 in ST-1901 was generated from *penA*-10 via recombination with another *Neisseria* species, followed by a second recombination event with a gonococcal strain harboring wildtype *penA*-1. Following the acquisition of *penA*-10 in ST-7363, a dominant sub-lineage rapidly acquired fluoroquinolone resistance mutations at GyrA 95 and ParC 87-88, possibly due to independent mutations rather than horizontal gene transfer. Literature data suggest the emergence of these resistance determinants may reflect selection from the standard treatment regimens in Japan at that time. Our findings highlight how recombination and antibiotic use across and within *Neisseria* species intersect in driving the emergence and spread of drug-resistant gonorrhea.

**Author summary:** Antimicrobial resistance is recognized as one of the greatest threats to human health, and *Neisseria gonorrhoeae* resistance is classified as one of the most urgent. The two major internationally spreading lineages resistant. to first line drugs likely originated in Japan, but when and how their genetic resistance determinants emerged remain unknown. In this study, we conducted an evolutionary analysis using clinical *N. gonorrhoeae* isolates from 37 countries, including a historical collection of Japanese isolates, to investigate the emergence of resistance in each of the two major lineages. We showed that the *penA* allele responsible for resistance to cephalosporins, the first-line treatment for gonorrhea, was possibly generated by two recombination events, one from another *Neisseria* species and one from another *N. gonorrhoeae* lineage. We also showed that mutations responsible for resistance to a previously widely used antibiotic treatment occurred twice independently in one of the two major lineages. The emergence of the genetic resistance determinants potentially reflects selection from the standard treatment regimen at that time. Our findings highlight how recombination (horizontal gene transfer) and antibiotic use across and within a bacterial species intersect in driving the emergence and spread of antimicrobial resistance genes and mutations.

## Introduction

Antimicrobial resistance (AMR) is one of the greatest threats to human health, urgently requiring effective global surveillance [1]. Gonorrhea is one of the most common sexually transmitted bacterial infections worldwide causing substantial morbidity and economic loss [2–4], with antimicrobial resistance an increasing concern. Among antimicrobial resistance mechanisms, *Neisseria gonorrhoeae* resistance to third-generation cephalosporins (3GCs) and fluoroquinolones has been defined as “high” priority by the World Health Organization (WHO). Among 3GCs, cefixime is no longer recommended for single-dose treatment, leaving ceftriaxone as the only remaining option for empirical first-line monotherapy in most countries [5]. Reduced susceptibility to cefixime and ceftriaxone is mainly caused by mutations in *penA*, the gene encoding the 3GC target, penicillin-binding protein 2 (PBP2). Mosaic *penA* alleles are deemed ‘mosaic,’ as the 3′ segments of their DNA sequences have been imported from other *Neisseria spp.* via homologous recombination events [6]. Many mosaic alleles have been documented, reflecting both distinct recombination events and additional mutations [7], and many of these mosaic alleles increase resistance to 3GCs.

Two internationally-disseminated sequence types (STs), defined by *Neisseria* multilocus sequence typing (MLST), ST-7363 and ST-1901, have acquired mosaic *penA* alleles and can be resistant to 3GCs. ST-1901, harboring mosaic *penA*-34, accounted for the majority of isolates with reduced susceptibility to 3GCs in the USA and Europe from the 2000s to at least the early 2010s [8, 9]. In Japan, ST-7363 accounted for the highest proportion (66%) among 149 isolates with reduced susceptibility to cefixime, isolated from 1998 to 2005 [10], and for 21% cases among 90 isolates obtained in 2015 [11]. These two STs harbor the mosaic *penA* alleles 34 and 10, which have identical nucleotide and amino acid sequences, except for those at the C-terminus (encoding 33 amino acids, approximately 6% of the entire sequence) [7, 11]. Both STs likely originated in Japan [12]. Recently, we analyzed whole genome sequence data combined with antimicrobial susceptibility testing results from 204 isolates from genomic surveillance in 2015 of a region where the first extensively drug resistant (XDR) *N. gonorrhoeae* resistant to ceftriaxone was isolated. This analysis was complemented with data from 67 further genomes from other time frames (from 1996 to 2015) and locations within Japan [11]. We first clustered STs that were closely related at the core-genome level to ST-1901 and ST-7363, resulting in ST-1901-associated and ST-7363-associated core genome groups. We found distinct evolutionary pathways of mosaic *penA* acquisition for these two core-genome groups: the ST-7363-associated core-genome group acquired *penA*-10 once, whereas the ST-1901-associated core-genome group had multiple independent acquisitions of *penA*-10 and *penA*-34. The previous study analyzed only Japanese *N. gonorrhoeae* isolates; thus, when and how the mosaic *penA* alleles—particularly *penA*-34, which is now dominant in USA and Europe—spread to other countries remained unknown. In addition, the majority of previously analyzed isolates were from 2015, and inclusion of additional genomes from other time-frames did not provide sufficient resolution to identify the time of emergence of the mosaic *penA* alleles with a narrow credibility interval [11]. Although the generation of *penA*-10 was explained by a single horizontal gene transfer (HGT) or recombination event, which imported the 3′ *penA* segment from a commensal *Neisseria spp.*, whether generation of *penA*-34 is similarly explained by a single HGT event or by multiple successive events remained unknown. Furthermore, the distribution of fluoroquinolone-resistance determinants, i.e., mutations in *gyrA* and *parC* [9], was examined only in strains isolated in 2015; however, when and how those mutations emerged and spread was also unclear.

In the present study, we addressed these issues by (i) obtaining complete, closed, genome sequences of three Japanese ST-1901 *N. gonorrhoeae* isolates harboring *penA*-34 from 2005, and (ii) reconstructing a dated phylogeny based on core-genome alignment of 1,075 genome sequences from isolates sampled from 37 countries. We also sequenced the genomes of 103 Japanese ST-7363 *N. gonorrhoeae* isolates from 1996–2005 and reconstructed a dated phylogeny of these genomes and 88 previously-sequenced Japanese ST-7363 genomes dating from 1996 to 2015. We further examined how these estimated dates corresponded to contemporary antibiotic use and treatment regimens. This generated a detailed narrative of the emergence and evolution of the cephalosporin-resistance genes and fluoroquinolone-resistance mutations, based on the dated phylogenies of these two global prevalent STs. These observations improve our understanding of how resistance determinants evolve via interactions among lineages and species under the selective pressure of antibiotic use.

## Results

### ST-1901-associated core-genome group and resistance to the 3GCs

A clonal dated phylogeny of ST-1901-associated core-genome group gonococci with branch lengths corrected to account for homologous recombination was inferred using ClonalFrameML, followed by BactDating (Fig 1) for 1,075 isolates sampled from 37 countries dating from 1992 to 2016 (S1 Table). Identified core-genome groups corresponded to the recently designated “core-genome group cluster 3”, which had been identified using a locus threshold of 400 or fewer locus differences [13] (S1 Table), except for 2 isolates, which belonged to cgc_400 groups 18 and 221. Parameter estimates from ClonalFrameML were consistent with those observed previously [11] (Supplementary text). In general, isolates harboring the mosaic *penA* alleles (colored cyan in Fig 1) and others were separated into two clusters, hereafter named ‘sub-lineage 34’ and ‘susceptible sub-lineage’, respectively. The sub-lineage 34 accounts for the majority of isolates encoding mosaic *penA* alleles, and harbors *penA-*34 and its variants (highlighted in the 1^st^ column titled “sub-lineage 34” of Fig 1 in yellow). This lineage was estimated to have emerged (red circle in the tree) between May 1990 and July 1999 (95% credibility interval). The sub-lineage 34 is shown as an enlargement in addition to its phylogenetic neighbor located at the bottom in S1 Fig. As shown in the 2^nd^ column headed “continent” in Fig 1, most strains were isolated from North America (light pink, 50.6%) and Europe (pink, 42.6%) mainly through the Gonococcal Isolate Surveillance Project (GISP) [8] (accounting for 80.1% of the strains isolated from North America) and the European Gonococcal Antimicrobial Surveillance Programme (Euro-GASP) [14] (accounting for 64.3% of the strains isolated from Europe), respectively, while very few strains were isolated from Asia (blue, 3.7%), Oceania (brown, 2.3%), and South America (light blue, 0.9%)In the sub-lineage 34, strains with the oldest isolation date were from 2005, among which 3 were from Japan and 2 from the USA. Complete genome sequences of the three Japanese strains were phylogenetically located at the base of the sub-lineage 34 tree (highlighted in the 3^rd^ column “complete genome” of Fig 1 in blue). The three Japanese isolates (GU029, GU058, and GU092) were collected in the Japanese prefectures of Aichi and Gifu in February–March 2005, indicating that none of them was likely closely related to an ancestor of the strains that spread to other countries.

**Fig 1.**
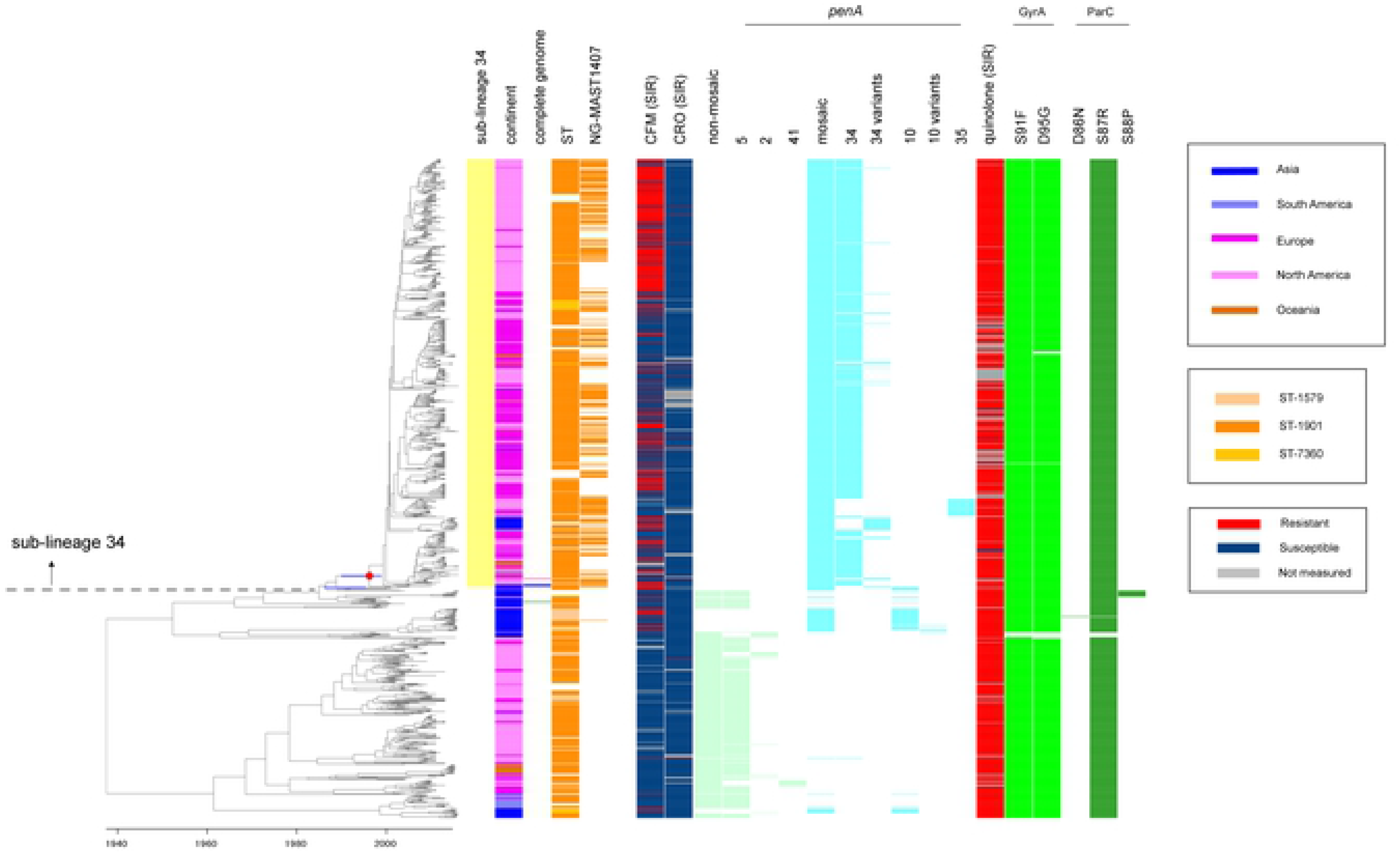
Whole genome sequence dated phylogeny, resistance patterns of the antimicrobials, and genetic polymorphisms in the ST-1901-associated core-genome group. Left: A clonal dated phylogeny with corrected branch lengths to account for homologous recombination. In the heat map, the 1^st^ column titled “sub-lineage 34”, the sub-lineage harboring *penA*-34 is colored. The 2^nd^ column titled “continent” shows Asia (deep blue), South America (light blue), Europe (pink), North America (light pink), or Oceania (brown). The 3^rd^ column titled “complete genome” shows the three Japanese strains harboring *penA*-34 from 2005 in blue and the ancestral Japanese strain (at the bottom) harboring *penA*-5 in black, whereas the WHO_Y (F89) strain is in pink. In the 4^th^ column titled “ST”, ST-1901 and its single locus-variants are colored using different colors as shown in our previous study [11]. In the 5^th^ column titled “NG-MAST1407”, strains of NG-MAST1407 are colored. In the 6^th^ and 7^th^ column, susceptible/resistant (S/R) categories according to the EUCAST minimal inhibitory concentration (MIC) breakpoint (susceptibility ≤ 0.125 μg/mL and resistance > 0.125 μg/mL) of 3GCs (cefixime CFM and ceftriaxone CRO) are shown. The columns were colored grey when the MIC values were missing. In the 8^th^ column, the presence (light yellow-green) or absence of any non-mosaic *penA* allele is shown. In the 9^th^–12^th^ columns, the presence (light yellow green) or absence of a specific non-mosaic *penA* allele is shown. The 13^th^ column shows the presence (cyan) or absence any mosaic *penA* allele. The 14^th^–18^th^ columns show presence (cyan) or absence a specific mosaic *penA* allele (specifically, 34 and its variants [54], 10 and its variants [54], and 35 [22]). The 19^th^ column shows the susceptible/resistant (S/R) categories of fluoroquinolones (mostly ciprofloxacin, and much less frequently, levofloxacin) according to the EUCAST breakpoint. The 20^th^–21^st^ columns show the presence (yellow-green) or absence of nonsynonymous amino acid changes compared to the wild type in GyrA. The 22^nd^–24^th^ columns show the presence (green) or absence of nonsynonymous amino acid changes compared to the wild type in ParC. In the clonal dated phylogeny at the left, a red circle indicates emergence of the sub-lineage 34, whereas two purple lines indicate 95% confidence intervals examined in the main text (emergence time of the sub-lineage 34, and that of one of the three sub-lineages harboring *penA*-10).

To provide an outgroup for sub-lineage 34, we obtained the complete genome sequence of an isolate, DRR129099 (12-032 in our previous study [11]), from the Japanese prefecture of Kanagawa in July 2000. This isolate harbors a non-mosaic *penA* allele and is phylogenetically close to the sub-lineage 34 (highlighted in green in the 3^rd^ column headed “complete genome” of Fig 1). Alignment of the complete genomes of the four isolates and the reference isolate WHO_Y (F89) included in the sub-lineage 34 (highlighted in pink in the 3^rd^ column headed “complete genome” of Fig 1), revealed conserved genomic synteny among the reference isolate in the sub-lineage 34, DRR129099 harboring a non-mosaic *penA*-5 allele, and one of the isolates (GU092) encoding *penA*-34 (1^st^, 4^th^, and 5^th^ genome in S2 Fig). This indicated that the overall genomic structure was maintained during evolution from the ancestor of the isolates in the sub-lineage 34 that spread to other countries.

In the whole ST-1901-associated core-genome group, ST-1901 and its single locus variants, ST7360 and ST1579, are colored in the “ST” column in Fig 1. The 91 (8.5%) other strains comprised 21 STs. The top five STs (ST-9365, ST-10312, ST-8153, ST-10241, and ST-13840) accounted for 62.6% of the samples (S3 Fig) and belonged to the ST-1901-associated core-genome group, although the nucleotide sequences of the seven loci differed from those of ST-1901. Similarly, in the sub-lineage 34, 51.7% of the strains were NG-MAST1407 (colored in the “NG-MAST1407” column in Fig 1), while the other strains were classified into 274 types or as undetermined according to the database of nucleotide sequences of the two highly variable NG-MAST loci (*porB* and *tbpB*).

Most isolates harboring the ‘non-mosaic’ *penA*-5 were located in the susceptible sub-lineage, whereas other isolates harboring *penA*-5 were present immediately outside the sub-lineage 34, suggesting that *penA*-5 was ancestral and that *penA*-34 evolved from it (Fig 1).

The sub-lineage harboring *penA*-34 and its variants appeared to be monophyletic, whereas the entire phylogeny of ST-1901-associated core-genome group included three separate mosaic *penA*-10 allele acquisitions, one of which was observed in the present study to be phylogenetically close to sub-lineage 34. This included the oldest isolates harboring *penA*-10 from 2000–2001 in Japan [15] (S1 Fig), which were not included in the previous study [11]. This suggests that *penA*-34 was generated via *penA*-10 from the ancestral *penA*-5 allele.

A small cluster of isolates encoding mosaic *penA*-35 was found in the sub-lineage 34, which consisted of 27 isolates that were isolated in India, Canada, UK, and USA from 2008 to 2011, and were not resistant to cefixime and ceftriaxone. A comparison of the amino acid sequences of *penA*-34 and 35 indicated that the differences in the amino acid residues across the sequence were likely consequences of HGT events spanning the entire *penA* locus from an unknown source outside the sub-lineage 34 (S4 Fig).

### Dated phylogeny of ST-7363-associated core-genome group, susceptibility to the 3GCs, and *penA* alleles

A clonal dated phylogeny of ST-7363-associated core-genome group, corresponding to recently designated “core-genome group cluster 8” [13] (Fig 2), showed that the isolates broadly separated into two clusters, corresponding to the presence or absence of a mosaic *penA* allele. In contrast to the ST-1901-associated core-genome group, we observed that the majority of the ancestral non-mosaic *penA* allele was *penA*-2, the nucleotide sequence of which is identical to that of *penA*-5 except for the region from nucleotides 1598–1752, the 3’ terminus of the coding sequence (*penA-*2 was also referred to as “*penA*-5 variant” [11]). Consistent with the results of our previous report, the majority of the mosaic *penA* was *penA*-10, with the exception of isolate H041 which was resistant to ceftriaxone [7], and one isolate encoding *penA*-34 (ERS311596, isolated in 2001) [15]. The phylogeny showed that *penA*-10 in ST-7363 was generated from the ancestral *penA*-2 (Fig 2) (95% credibility interval September 1991–May 1995). In agreement with a prior observation [11], we noted that *penA*-150 (designated in the NG-STAR database, Fig 2), susceptible to 3GCs, appeared to be a product of the recombination between the mosaic *penA*-10 or 37 (H041-type) alleles and *penA*-5.

**Fig 2.**
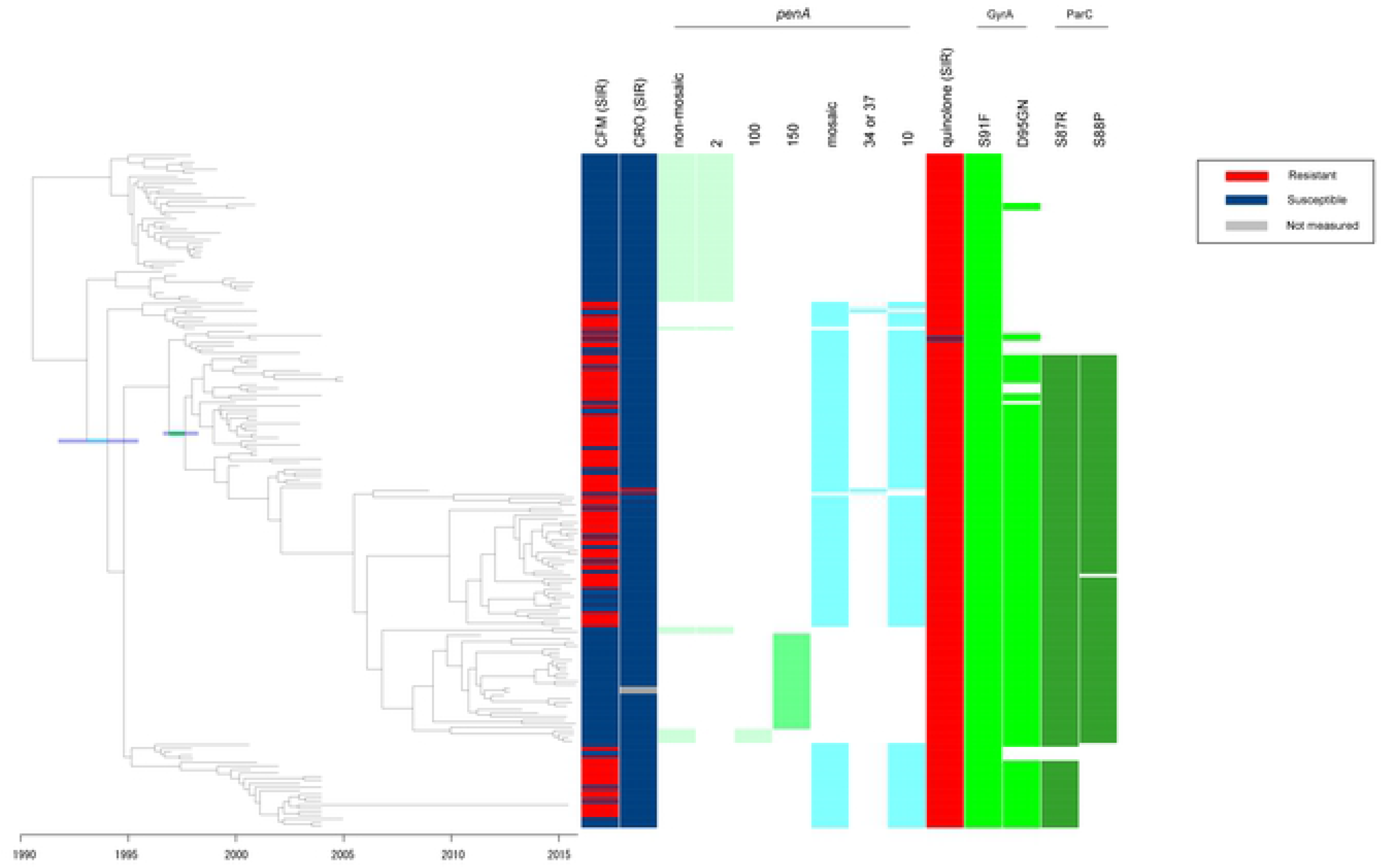
Whole genome sequence dated phylogeny, resistance patterns of the antimicrobials, and genetic polymorphisms in the ST-7363-associated core-genome group. The columns in the heat map are almost the same as those in Fig 1, although the first five columns in Fig 1 have been omitted here. The 1^st^ and 2^nd^ columns show the susceptible/resistant (S/R) categories according to the EUCAST breakpoint of cefixime (CFM) and ceftriaxone (CRO). The 3^rd^ column shows the presence (light yellow-green) or absence of any non-mosaic *penA* allele. The 4^th^–6^th^ columns show the presence (light yellow-green) or absence of a specific non-mosaic *penA* allele. The 7^th^ column shows the presence (cyan) or absence of any mosaic *penA* allele. The 8^th^–9^th^ columns show the presence (cyan) or absence a specific mosaic *penA* allele (specifically, 10, and 34 or 37 (H041-type)[7]). The 10^th^ column shows the susceptible/resistant (S/R) categories of fluoroquinolones (mostly ciprofloxacin, and much less frequently, levofloxacin) according to the EUCAST breakpoint. The 11^th^–12^th^ columns show the presence (yellow green) or absence of nonsynonymous amino acid changes compared to the wild type in GyrA. The 13^th^–14^th^ columns show the presence (green) or absence of nonsynonymous amino acid changes compared to the wild type in ParC. In the clonal dated phylogeny at the left, the two branches of interest examined in the main text are colored cyan and green, with 95% confidence interval of the two evolutionary events (acquisition of *penA*-10 and simultaneous amino acid substitutions at GyrA 95 and ParC 87–88) colored purple.

### Proposed origin of mosaic *penA*-34 and 10 alleles

To further analyze the origins of mosaic *penA*-34 and *penA*-10, we compared their nucleotide sequences to those of the susceptible alleles in the ST-1901 and ST-7363-associated core-genome groups. Nucleotide sequence alignment of *penA*-5, 10, and 34 and their downstream sequences in ST-1901 is shown schematically in Fig 3, and at the nucleotide level in S5 Fig. The 1^st^ to 293^rd^ nucleotides (region (1) in Fig 3A) were identical, followed by the recombined mosaic region from 294^th^ nucleotide (region (2), indicated by the red line in *penA*-10 in Fig 3A, and between red arrows in S5 Fig), in which sequence identity between *penA*-5 and *penA*-10 with their downstream sequences was 89.0%, whereas sequence identity between *penA*-10 and *penA*-34 with their downstream sequences was 98.2%. The coding sequences (CDS) of *penA-*10 and 34 were identical, with the exception of 105 bp at the 3’ end, indicated by the left part of the orange line at the end of *penA*-34 in Fig 3A. In the following region (3), the downstream sequences of *penA*-5 and *penA*-10 were identical, whereas those of *penA*-10 and 34 harbored four individual polymorphisms and 99.6% sequence identity. In the following region (4), the nucleotide sequences of *penA*-5, 10, and 34 exhibited ≥ 99.9% sequence identity. Based on the results of nucleotide sequence comparison, the origin of *penA*-10 was most parsimoniously explained by an approximately 2.4 kb recombination event from the start of mosaic region (2) in a *penA*-5 background. The source is unknown, as BLASTn search against the NCBI nr database did not yield any hit in other *Neisseria* species with high (> 95%) sequence identity.

**Fig 3.**
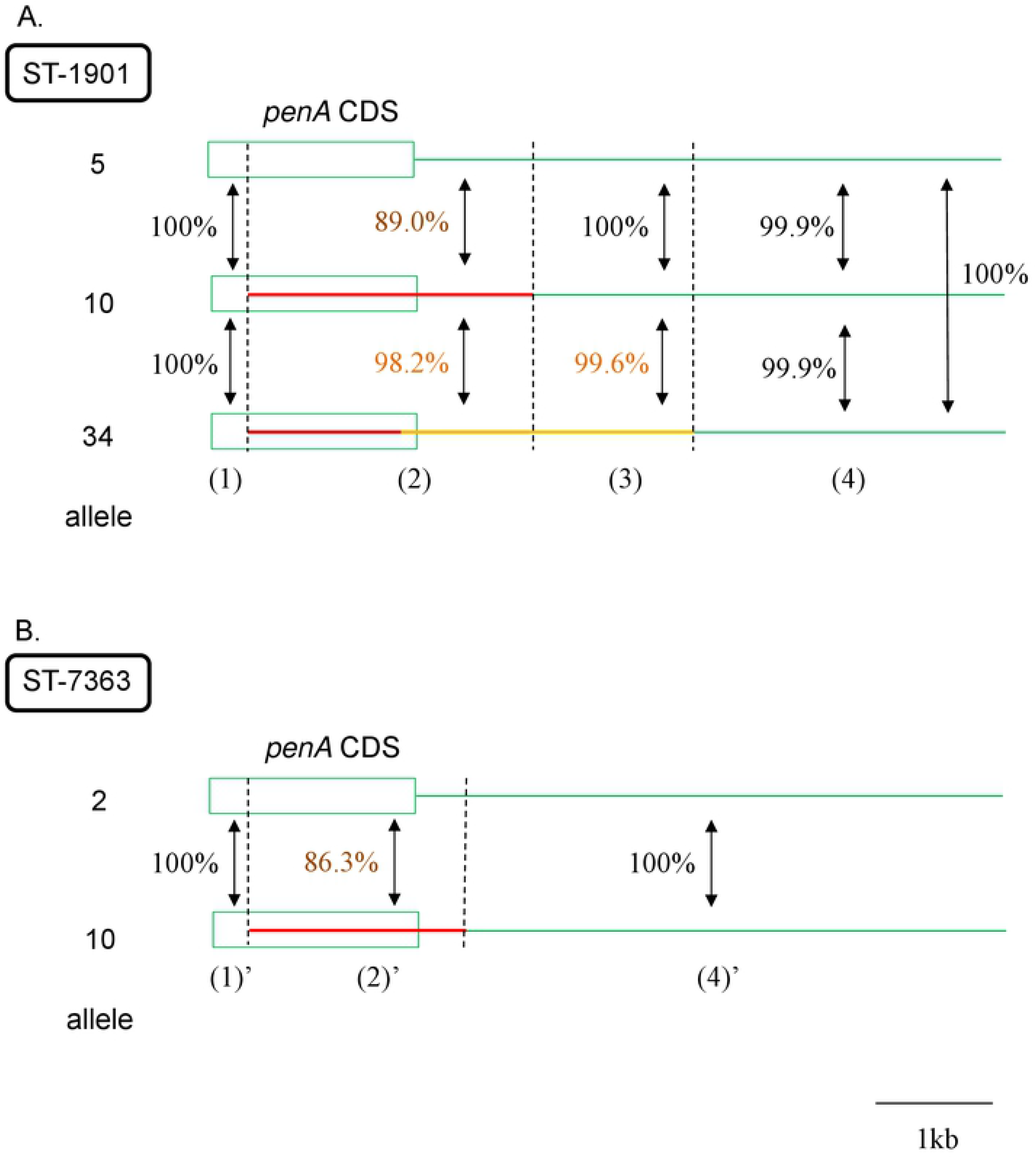
Schematic depiction of nucleotide sequence alignment of *penA* and its downstream sequence. (A) ST-1901-associated core-genome group. *penA*-5 (top), 10 (middle), and 34 (bottom). (B) ST-7363-associated core-genome group. *penA*-2 (top) and 10 (bottom). The coding sequence (CDS) of *penA* is shown as a rectangle. The recombined sequences in *penA*-10 and 34 are indicated by the red and orange lines, respectively.

PenA 34 and 10 differed in the C-terminus, and the difference continued in their downstream nucleotide sequences in the regions (2) and (3). To investigate the possibility that *penA-*34 emerged via HGT in the background of *penA*-10, we aligned the nucleotide sequence corresponding to the orange line in regions (2) and (3) (approximately 2.5 kb, between orange arrows in S5 Fig) to the genome sequences of 140 strains isolated in 1996–1997 in the Japanese prefecture of Kanagawa (S3 Table, most of them newly sequenced in the present study) and the 204 Japanese strains isolated in 2015 [11], representing multiple STs. A maximum likelihood tree constructed from this alignment (S7 Fig) revealed that ST-1901 isolates encoding *penA*-34 and its variant were clustered with ST-1594 strains encoding *penA*-1. In particular, the nucleotide sequence was identical between *penA*-34 and *penA*-1, with the exception of a single polymorphism whereas the *penA*-34 variant harbored a polymorphism at the C-terminus. The next most similar sequence was found in a strain (09-021) harboring *penA-* 2 (located next to the ST-1594 strains encoding *penA*-1 in S7 Fig) and containing five polymorphisms compared to the sequence of *penA*-1. These results suggested that the origin of *penA*-34 can be explained by two recombination events: first, a recombination event from a commensal *Neisseria* into a *penA*-5 background, resulting in *penA*-10 (red line in *penA*-10 in Fig 3A); second, a recombination event from *penA*-1 into the *penA*-10 background resulting in *penA*-34 (orange line in *penA*-34 in Fig 3A and orange rectangle in Fig 4).

**Fig 4.**
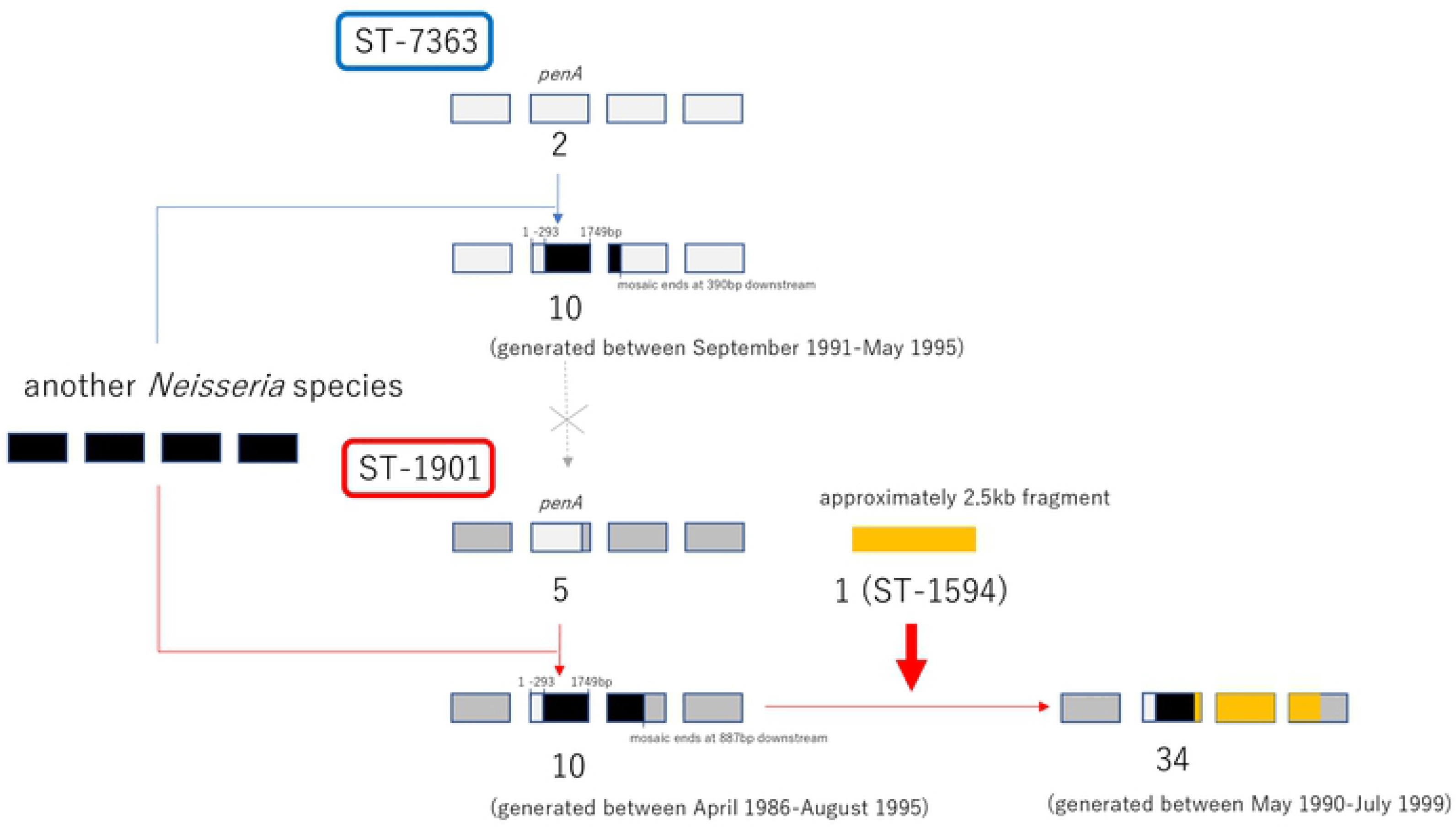
Schematic depiction summarizing the results regarding the origin of mosaic *penA*-10 and 34. Rectangles indicate genes, second of which is *penA*. In ST-7363 (top) and ST-1901 (bottom), the upper part shows the ancestral sequences (*penA*-2 and 5, respectively) while the lower part shows the recombined sequences.

Although no study has reported the prevalence of the *penA*-1 allele in Japan, analysis of genome sequence data of the 140 isolates in 1996–1997 and those of 204 isolates in 2015 collected via surveillance of symptomatic cases, showed that the *penA*-1 allele was encoded only in ST-1594 that accounted for 3.6% and 7.8% cases (the 7^th^ and 5^th^ most frequent ST) among the collected strains in 1996–1997 and 2015, respectively.

Similar to ST-1901, *penA*-2 and 10 of ST-7363 exhibited identical nucleotide sequence from nucleotide 1 to 293, followed by the mosaic region from nucleotide 294 to 390 bp downstream of the end of *penA* (region (2)’, indicated by the red line in Fig 3B and between red arrows in S6 Fig). This is most parsimoniously explained by a recombination event, although the source of this DNA segment is unknown (Fig 4). Comparison of the length of the recombined regions between ST-1901 and ST-7363 indicated that *penA*-10 in ST-7363 was unlikely to be the origin of *penA*-10 in ST-1901, as the recombined fragment was shorter than that of ST-1901 (Fig 4).

### Analysis of susceptibility to fluoroquinolones and examination of mutations in *gyrA* and *parC* in the dated phylogenies

For fluoroquinolone resistance, the dated phylogenies also revealed distinct evolutionary paths for the ST-1901- and ST-7363-associated core-genome groups. Almost all isolates in both core-genome groups were resistant to fluoroquinolones (minimal inhibitory concentration (MIC) > 0.06 μg/mL, using the EUCAST breakpoint) and harbored the GyrA 91F substitution. In the ST-1901-associated core-genome group, the substitutions at GyrA D95G and ParC S87R were shared among most isolates, including the clade encoding the non-mosaic *penA* (Fig 1). The substitution at ParC S88P was found only in a small cluster of 11 strains (bottom of Fig 1). In contrast, in the ST-7363-associated core-genome group, a dominant sub-lineage contained amino acid substitutions at GyrA D95GN, ParC S87R, and ParC S88P, which arose within a short time (highlighted as a green horizontal line in Fig 2; 95% credibility interval August 1996–March 1998). These substitutions arose after the acquisition of *penA*-10, which is inferred to have occurred on a more ancestral node: the 95% credibility interval for the dates of these two events do not overlap. The output of ClonalFrameML used for constructing the clonal phylogeny did not include a signature of the recombination importing the GyrA and ParC substitutions, suggesting that they arose via independent mutations.

## Discussion

An extensive analysis of *N. gonorrhoeae* ST-1901 and ST-7363-associated core-genome groups, including the 1,075 isolates sampled from 37 countries for ST-1901 and 191 Japanese isolates sampled for ST-7363, has improved the resolution of the time-resolved phylogenies of these important gonococci. Newly sequenced, historical, Japanese isolates included three ST-1901 gonococci harboring the mosaic *penA*-34 (NG-MAST1407), which accounted for the majority of isolates with reduced susceptibility to 3GC in the USA and Europe. In the ST-1901-associated core-genome group, the three earliest isolates were obtained in Japan from February–March, 2005, while two originated in the USA (GCGS0944, GCGS0920) in 2005 (S1 Fig). The time-resolved tree suggested that the three Japanese isolates were phylogenetically closest to the unsampled ancestor, consistent with the hypothesis that this sub-lineage likely originated in Japan [12].

The time-resolved trees of ST-1901 and ST-7363-associated core-genome groups enabled us to estimate the timing of the emergence of three genetic determinants of antimicrobial resistance: (i) mosaic *penA*-34 in ST-1901 (May 1990–July 1999); (ii) mosaic *penA*-10 in ST-1901 (April 1986–August 1995) and in ST-7363 (September 1991–May 1995); and (iii) GyrA D95GN and ParC S87R-S88P in ST-7363 (August 1996–March 1998).

How do these estimated dates correlate to antibiotic use? Although there were no national data on antibiotic usage at that time in Japan, a 1999 survey on the use of antimicrobial drugs in 14 hospitals and 4 clinics mostly in a prefecture in Japan showed that 50% cases (9 out of 18) exclusively used fluoroquinolones for the treatment of urethritis (Dr. Mitsuru Yasuda, unpublished data). Similarly, another survey of clinics in Fukuoka in Japan in the late 1990s reported that “Fluoroquinolones were most frequently used (46%) as first-line treatment for gonorrhea, followed by penicillin with or without a β-lactamase inhibitor (28%), cephems (16%), and others (10%). Of the various fluoroquinolones available in Japan, a 7-day course of levofloxacin (200 mg twice a day or 100 mg three times a day) was most frequently used in the treatment of gonococcal infections (M. Tanaka, unpublished data)” [16]. The regimen used in Japan in the late 1990s exceeded the single ciprofloxacin dose recommended in the USA, and may have contributed to selection of the *gyrA* and *parC* mutations, resulting in increased fluoroquinolone resistance.

A survey in Fukuoka, Japan, also showed that in the 1990s, fluoroquinolones were most frequently used (46%) as the first-line of treatment for gonorrhea, followed by penicillin with or without a β-lactamase inhibitor (28%), and cephems (16%) [16]. In 1999, the reported MIC_90_ for cefixime for clinical isolates of *N. gonorrhoeae* was 0.06 to 0.125 μg/mL, which indicated the presence of isolates with cefixime MIC = 0.125 μg/mL. This prompted the conclusion that a single dose of 400 mg cefixime used in the USA would not eradicate ≤ 95% of the clinical strains of *N. gonorrhoeae* [17]. The first Japanese STI treatment guidelines in 1999 described a regimen of six 100 mg doses of cefixime at a 12-h interval. Between 1999 and 2001, a regimen of 200 mg cefixime at 6 h interval was proposed, although it eradicated only 88.2% and 54.5% *N. gonorrhoeae* isolates with MIC_90_ and MIC values of 0.125 μg/mL, respectively [17]. These regimens and the observed circulating resistance might have contributed to the selection and dissemination of gonococcal strains with reduced susceptibility to cephalosporins. Although information regarding the alleles responsible for the cefixime MIC of 0.125 μg/mL at that time is lacking, the rising MICs and shifting patterns of antibiotic use reflect the dynamic interplay between the evolution of cefixime resistance and shifting treatment strategies.

Regarding the GyrA D95GN and ParC S88P amino acid substitutions in ST-7363, responsible for fluoroquinolone resistance, it is interesting to note that a sub-lineage (Fig 2) harboring two substitutions (GyrA D95GN and ParC S87R), but not at ParC 88, has not been sampled since 2005 (except for the GU294 strain isolated in 2015 as indicated by the long branch at the bottom of Fig 2). The apparent loss of this sub-lineage suggests the importance of the substitution at ParC 88 or another mutation that became dominant in ST-7363. Alternatively, the stochasticity of transmission or some other factor (perhaps susceptibility to local treatments) might have led to the loss of this sub-lineage.

We propose two potential recombination events for the generation of *penA-*34 in ST-1901: recombination with another *Neisseria* species that generated *penA*-10, followed by another recombination event with a strain harboring *penA*-1 that converted *penA*-10 to *penA*-34. The lengths of the recombined sequence were approximately 2.3 kb and 2.5 kb, respectively, which is consistent with a recent estimate (2.5 kb) of the mean of the geometrically distributed DNA tract lengths transferred between donors and recipients in *N. gonorrhoeae* [18]. Similarly, recombination with a donor susceptible to 3GC was recently reported in *N. gonorrhoeae* in Japan [11] and the USA [8, 19], although it led to loss of the resistance phenotype. It should be noted that the inference of recombination is based on sampling, and we cannot rule out the possibility that recombination with a currently unsampled strain might have generated *penA*-34. Nevertheless, we can speculate that a strain harboring *penA*-1 is currently the most likely donor of the recombination that generated *penA*-34 from *penA-*10.

The reason behind the dominance of ST-1901 harboring *penA*-34, but not ST-7363 harboring the same allele, remains unsolved. In the ST-7363 group, *penA*-10 was dominant, and there was only one isolate with *penA*-34, identified in 2001. Data indicating significant difference in fitness between *penA*-10 and 34 in vitro are lacking. Possibly, the differences in fitness, selection, and dissemination between *penA*-10 and 34 depended on the genetic background of those sequence types and environmental factors at the time they were selected.

Unfortunately, data regarding the potential environmental factors such as national or regional information of antimicrobial use at the time of their emergence and dissemination are not accessible. Further studies are therefore warranted to prospectively collect such data and isolates, and conduct integrative analyses of genome sequences, antimicrobial susceptibility, and the environmental factors in order to monitor, understand, and control emergence and dissemination of new resistance determinants of public health importance. A recent study that analyzed the genome and antimicrobial susceptibility data of *N. gonorrhoeae* isolates from 1928 to 2013 in Denmark [20], reported that no isolate was interpreted to be ciprofloxacin-resistant (MIC > 0.06 μg/mL) until the 1980s. The percentage of isolates resistant to ciprofloxacin increased from 0 to 14.3% in 1990s. Five isolates from the 1950s–1970s contained a GyrA S91T amino acid substitution, although all of them were susceptible to ciprofloxacin. Compared to the recent study, the isolates analyzed in our study were all collected after 1990: the oldest strain of the ST-1901-associated core-genome group was collected in 1992, and that of ST-7363 in 1996. Further studies are warranted to conduct genome sequencing of historical Japanese strains collected before 1990s to identify isolates phenotypically susceptible to fluoroquinolones and explore when and how the individual amino acid substitutions of GyrA S91F, D95G, and ParC S87R in ST-1901 and that of GyrA S91F in ST-7363 may have occurred.

Our previous study revealed that the different evolutionary pathways of the two major core-genome groups regarding the mosaic *penA* alleles responsible for resistance to 3GC [11]. The present study increased our understanding by elucidating the following three major points: 1) *penA*-34 in ST-1901 was likely generated from *penA*-10 via two recombination events; 2) the *penA*-10 allele in the ST-1901-associated core-genome group emerged in at least three distinct Japanese sub-lineages independently, one of which was phylogenetically adjacent to the sub-lineage harboring *penA*-34; and, 3) the *penA*-10 allele in ST-7363, possibly generated via recombination with another *Neisseria* species, was unlikely to be a source of HGT that generated *penA*-10 in ST-1901-associated core-genome group, as the recombined fragment was shorter than that of ST-1901-associated core-genome group. The single acquisition of *penA*-10 in ST-7363 was revealed in both the previous and present studies, although our understanding regarding its relationship with *penA*-10 and *penA*-34 in the ST-1901-associated core-genome group in terms of order of their generation and possibility of recombination between the two core-genome groups were improved in the present study. Furthermore, the present study demonstrated another interesting difference in fluoroquinolone resistance between the two core-genome groups since 1990s; ST-7363 was originally susceptible to 3GC and harbored amino acid substitution only in GyrA 91 in the 1990s; the simultaneous amino acid substitutions at GyrA 95 and ParC 87 occurred in the sub-lineage in the short time after it acquired the mosaic *penA*-10, whereas in the ST-1901-associated core-genome group, all the substitutions in GyrA 91, 95, and ParC 87 were already observed in 1990s in most strains (99.2%, 1066/1075), including the sub-lineage mostly susceptible to 3GC (at the bottom in Fig 1).

In summary, after combining the previously published dataset from 37 countries with the new genome sequence data and the antimicrobial susceptibility data of historical gonococcal isolates from Japan, we described the possible pathways of emergence of cephalosporin-resistant genes and fluoroquinolone-resistant mutations of two globally circulating *N. gonorrhoeae* core-genome groups. We further discussed the dynamic interplay between the evolution of antibiotic resistance and treatment regimens during time period of the emergence of genetic determinants of antimicrobial resistance. Such elucidation of evolutionary pathways will be useful for understanding and controlling the current and future evolution and spread of the pathogen, and resistance determinants driven by recombination and selective pressure of antibiotic use.

## Methods

### Isolates and DNA sequencing

In total, 1,075 genome sequences from *N. gonorrhoeae* isolates belonging to a ST-1901-associated core-genome group were included (S1 Table). This also comprised three mosaic *penA-*34-harboring ST-1901 isolates from Japan (2 isolates from Aichi and 1 isolate from Gifu prefectures) isolated in February–March 2005 and newly sequenced in this study using MinION and MiniSeq. These three strains were isolated via surveillance by Gifu University that obtained 51 ST-1901 strains (95 strains across various STs) in 2005. In addition, a ST-1901 strain (12-032) isolated in 2000 in Kanagawa, Japan [11] harboring *penA*-5 and susceptible to cephalosporins was re-sequenced using the MinION platform to examine genomic synteny between it and the strains harboring *penA*-34. Five other ST-1901 isolates dating from 2001 to 2004 in Kanagawa, Japan, were sequenced using the Illumina MiSeq platform. These newly sequenced genomes were combined with those of 1,066 publicly available genomes from 14 studies covering 37 countries [8, 11, 14, 15, 19, 21–28]. The publicly available genomes were selected based on a tree constructed in our previous study [29] such that they formed a group at the core-genome level.

For MinION sequencing, a MagAttract HMW DNA Kit (Qiagen, Hilden, Germany) was used for isolation of high molecular weight genomic DNA of each isolate, and the Rapid Sequencing Kit (SQK-RAD004) and R9.4 flowcells were used. For MiSeq sequencing, the genomic DNA of each isolate was extracted using a MagNA Pure LC DNA isolation kit on a MagNA pure LC instrument (Roche Diagnostics GmbH, Mannheim, Germany), which was used for Nextera XT library construction and genome sequencing (300 bp paired-end) using an Illumina MiSeq Reagent Kit v3 (600-cycle).

For ST-7363, 103 strains that were isolated from 1996 and 2005 in Kanagawa were sequenced using the Illumina MiSeq platform. These data were combined with 88 publicly available genomes [11, 15] (S2 Table).

### Antimicrobial susceptibility testing

The MICs of 3GCs (ceftriaxone and cefixime) and fluoroquinolones (ciprofloxacin and levofloxacin) of the newly sequenced historical strains isolated in Kanagawa were determined using agar dilution method [30] at the Kanagawa Prefectural Institute of Public Health and Gifu University (S1 and S2 Tables). MICs measurements were repeated for the following strains for which the genotypes and phenotypes were initially discordant, and MICs that better matched the genotypes were subsequently used: GCGS0938 (ceftriaxone) and GCGS0627 (ceftriaxone) in ST-1901 using the Etest method (bioMérieux), and GU250 (cefixime, ceftriaxone), GU431 (cefixime, ceftriaxone), and GU478 (cefixime, ceftriaxone) in ST-7363 using the agar dilution method. To define susceptible/resistant phenotypes, the following MIC cut-offs were used according to the European Union Committee on Antimicrobial Susceptibility Testing (EUCAST; www.eucast.org/clinical_breakpoints): susceptibility ≤ 0.125 μg/mL and resistance > 0.125 μg/mL for ceftriaxone and cefixime; susceptibility ≤ 0.03 μg/mL and resistance ≥ 0.06 μg/mL for ciprofloxacin. For strains on which MICs of fluoroquinolones were measured only for levofloxacin by the Gifu University (with names starting by “GU_”), the MIC cut-off of ciprofloxacin was used, as that of levofloxacin is not defined in EUCAST and a previous study has shown that MIC_50_, MIC_90_, and MIC range of ciprofloxacin and levofloxacin were similar for a set of 87 isolates [31].

### Genome assembly

The Illumina read data (300 bp paired-end reads) of each isolate sequenced using MiSeq were used for de novo assembly using SPAdes [32], or Shovill [33]; SPAdes was used at its core, but speed was improved. The assembled contigs were checked to ensure that the coverage was more than 10× and that total genome size was approximately equal to that of the type strain *N. gonorrhoeae* FA1090 (2.0–2.3M bp). The MinION read data of each of the four isolates of ST-1901 were used for de novo assembly using Canu version 1.7 [34]. Errors were corrected using three runs of Pilon for each contig assembled from the MinION reads [35] based on mapping of Illumina reads to the contigs using bowtie2 [36] with “very-sensitive” option, followed by automated circularization of the contigs using Circlator [37] and adjustment of the 1^st^ position of the genome sequence to the start of *dnaA* using an in-house script. The number of contigs and N50 of each isolate are summarized in S1 and S2 Tables. All isolates used in this study can be found on the pubMLST.org/neisseria database where MLST, NG MAST and NG STAR STs were determined along with isolate core genome groups as described previously [13]. MLST typing of 158 out of the 1,075 strains did not result in ST-1901, but were included in our dataset, as they formed a large cluster with other ST-1901 strains as a core-genome group. Each genome was annotated using Prokka [38].

### Construction of the clonal time-resolved phylogeny of ST-1901-associated and ST-7363-associated core-genome groups

As in previous studies [11, 39], pairwise genome alignment between the reference genome and one of the other strains was performed using progressiveMauve [40] for SNP calling and evolutionary analyses in ST-1901 and ST-7363, which enabled the construction of positional homology alignments even for genomes with variable gene content and rearrangement. The complete genome sequences of the “WHO_Y” (F89) and “WHO_X” (H041) strains [41] were used as ST-1901 and ST-7363 references, respectively. The alignments were then combined into a multiple whole genome alignment, in which each position corresponded to that of the reference genome. This approach was validated [11] by comparing with another whole genome alignment using a pipeline [42] that similarly conducts pairwise genome alignment between a reference genome and one of the other genomes using MUMmer [43]. A maximum likelihood tree was constructed using PhyML [44] from the genome alignment containing the 10,844 core SNPs in ST-7363. A maximum likelihood tree was generated from the genome alignment containing the 28,306 core SNPs in ST-1901 using RAxML [45], as PhyML failed to complete the tree generating process. The default setting was used for RAxML, whereas for PhyML, we used the -m GTR -c 4 -a e parameters that indicate the GTR + G4 model of DNA substitution with estimation of the shape parameter of the gamma distribution by maximizing the likelihood. Using this as a starting tree, a clonal phylogeny was constructed with corrected branch lengths to account for homologous recombination using the standard model of ClonalFrameML [46]. The alignment sites that were present in at least 70% isolates were used to focus on core and fairly conserved sites with missing frequency ≤ 30%, as validated in our previous study [11]. TempEst [47] was used to root the phylogeny by minimizing the sum of the squared residuals from the regression of root-to-tips on sampling year. BactDating [48] was used, in which the significance of the temporal signal in both ST-7363 and ST-1901-associated core-genome groups was confirmed (the date randomization test, *P* < 10^−4^). BactDating was then run for 10 million Markov chain Monte Caro (MCMC) iterations. After completion, the effective sample size (ESS) of all parameters were confirmed as exceeding 400 and trace plots were confirmed as demonstrating well-mixed chains. A time-resolved phylogeny and 95% confidence interval for each branch in the tree was exported.

### Analysis of genetic AMR determinants

For 3GC resistance determinants, the *penA* alleles were extracted based on a BLASTn search of the nucleotide sequence of mosaic *penA* alleles-10 and-34 and that of non-mosaic *penA*-5 (https://figshare.com/articles/penA_nucleotide_sequence_alignment_penA_5_X_XXXIV_and_1_/12203858) against each of the other genomes. When the mosaic *penA* alleles extracted based on the BLASTn search were truncated, sequence reads were mapped to the nucleotide sequence of *penA* alleles-10 and-34. The nucleotide sequence identity between the alleles was checked, and the allele numbering defined in NG-STAR database (https://ngstar.canada.ca) [49] was assigned to each *penA* allele [“penA_allele (NG-STAR)” column in S1 and S2 Tables]. Fig 1 shows the grouping of “*penA*-34 variants” for alleles that were 100% aligned to *penA*-34 and showed sequence identity of < 100% and > 99.5%, which was higher than that observed with *penA*-10. The “*penA*-10 variants” were grouped similarly.

For fluoroquinolone resistance determinants, the nonsynonymous substitutions in *gyrA* and *parC* described in a recent review [9] and also examined in our previous study [11] were analyzed. In *N. gonorrhoeae*, quinolone resistance has been attributed to nonsynonymous substitutions in specific regions of *gyrA* and *parC*, namely, amino acid positions 91 and 95 in GyrA, position 87 in ParC, and less frequently positions 86 and 88 in ParC. The presence or absence of the substitutions was detected using PointFinder [50]. The presence or absence of these genetic AMR determinants, as well as the MICs of the antimicrobial drugs for each strain, were illustrated as heat maps using Phandango [51].

### Inference of recombination events that generated mosaic *penA*-10 and 34 alleles

To investigate the recombination events that generated the mosaic *penA* alleles, a nucleotide sequence alignment of *penA* and a 5 kb region downstream of its 3’ end was prepared based on a BLASTn search against the genome sequences of ST-1901-and ST-7363-associated core-genome groups. The nucleotide sequences of *penA*-34 and its downstream region were used as query sequences in the BLASTn search. The nucleotide sequences obtained from the BLASTn were aligned using MAFFT v7.245 [52] and manually examined using Jalview [53] to analyze nucleotide sequence identity and detect recombined fragments for *penA*-10 and 34 in ST-1901 and *penA*-10 in ST-7363. The BLASTn search was also performed against a custom database of genome sequences of Japanese *N. gonorrhoeae* strains: 140 strains isolated in 1996–1997 in the Japanese prefecture of Kanagawa (S3 Table) and 204 strains isolated via genomic surveillance in 2015 in the Japanese prefectures of Kyoto and Osaka [11], both of which were not confined to ST-1901 or ST-7363, but included various STs. The nucleotide sequences obtained using BLASTn were aligned using MAFFT v7.245 [52] and manually examined using Jalview [53], using which an alignment of the potentially recombined fragment was extracted. A maximum likelihood tree of the fragment was constructed using PhyML [44], which was mid-point rooted and visualized using FigTree version 1.4.3. To investigate the potential sources of the recombination, a BLASTn search was performed for each recombined fragment in the NCBI nt database or the custom database of genome sequences of Japanese *N. gonorrhoeae* strains.

## Acknowledgments

The computational calculations were performed at the Human Genome Center at the Institute of Medical Science (University of Tokyo) and at the National Institute of Genetics.

## References

1. Sugden R, Kelly R, Davies S. Combatting antimicrobial resistance globally. Nat Microbiol. 2016;1(10):16187. doi: 10.1038/nmicrobiol.2016.187. PubMed PMID: 27670123.

2. Newman L, Rowley J, Vander Hoorn S, Wijesooriya NS, Unemo M, Low N, et al. Global Estimates of the Prevalence and Incidence of Four Curable Sexually Transmitted Infections in 2012 Based on Systematic Review and Global Reporting. PLoS One. 2015;10(12):e0143304. doi: 10.1371/journal.pone.0143304. PubMed PMID: 26646541; PubMed Central PMCID: PMCPMC4672879.

3. WHO. Global incidence and prevalence of selected curable sexually transmitted infections 2008 [cited 2017 May 12]. Available from: http://www.who.int/reproductivehealth/publications/rtis/2008_STI_estimates.pdf.

4. WHO. Global action plan to control the spread and impact of antimicrobial resistance in Neisseria gonorrhoeae 2012 [cited 2017 May 12]. Available from: http://whqlibdoc.who.int/publications/2012/9789241503501_eng.pdf.

5. Whittles LK, White PJ, Didelot X. Estimating the fitness cost and benefit of cefixime resistance in *Neisseria gonorrhoeae* to inform prescription policy: A modelling study. PLoS Med. 2017;14(10):e1002416. doi: 10.1371/journal.pmed.1002416. PubMed PMID: 29088226; PubMed Central PMCID: PMCPMC5663337.

6. Bowler LD, Zhang QY, Riou JY, Spratt BG. Interspecies recombination between the penA genes of Neisseria meningitidis and commensal Neisseria species during the emergence of penicillin resistance in N. meningitidis: natural events and laboratory simulation. J Bacteriol. 1994;176(2):333–7. doi: 10.1128/jb.176.2.333-337.1994. PubMed PMID: 8288526; PubMed Central PMCID: PMCPMC205054.

7. Ohnishi M, Golparian D, Shimuta K, Saika T, Hoshina S, Iwasaku K, et al. Is *Neisseria gonorrhoeae* initiating a future era of untreatable gonorrhea?: detailed characterization of the first strain with high-level resistance to ceftriaxone. Antimicrob Agents Chemother. 2011;55(7):3538–45. doi: 10.1128/AAC.00325-11. PubMed PMID: 21576437; PubMed Central PMCID: PMCPMC3122416.

8. Grad YH, Harris SR, Kirkcaldy RD, Green AG, Marks DS, Bentley SD, et al. Genomic Epidemiology of Gonococcal Resistance to Extended-Spectrum Cephalosporins, Macrolides, and Fluoroquinolones in the United States, 2000-2013. J Infect Dis. 2016;214(10):1579–87. doi: 10.1093/infdis/jiw420. PubMed PMID: 27638945; PubMed Central PMCID: PMCPMC5091375.

9. Harrison OB, Clemence M, Dillard JP, Tang CM, Trees D, Grad YH, et al. Genomic analyses of *Neisseria gonorrhoeae* reveal an association of the gonococcal genetic island with antimicrobial resistance. J Infect. 2016;73(6):578–87. doi: 10.1016/j.jinf.2016.08.010. PubMed PMID: 27575582; PubMed Central PMCID: PMCPMC5127880.

10. Shimuta K, Watanabe Y, Nakayama S, Morita-Ishihara T, Kuroki T, Unemo M, et al. Emergence and evolution of internationally disseminated cephalosporin-resistant *Neisseria gonorrhoeae* clones from 1995 to 2005 in Japan. BMC Infect Dis. 2015;15:378. doi: 10.1186/s12879-015-1110-x. PubMed PMID: 26381611; PubMed Central PMCID: PMCPMC4574456.

11. Yahara K, Nakayama SI, Shimuta K, Lee KI, Morita M, Kawahata T, et al. Genomic surveillance of Neisseria gonorrhoeae to investigate the distribution and evolution of antimicrobial-resistance determinants and lineages. Microb Genom. 2018;4(8). doi: 10.1099/mgen.0.000205. PubMed PMID: 30063202; PubMed Central PMCID: PMCPMC6159555.

12. Unemo M, Nicholas RA. Emergence of multidrug-resistant, extensively drug-resistant and untreatable gonorrhea. Future Microbiol. 2012;7(12):1401–22. doi: 10.2217/fmb.12.117. PubMed PMID: 23231489; PubMed Central PMCID: PMCPMC3629839.

13. Harrison OB, Cehovin A, Skett J, Jolley KA, Massari P, Genco CA, et al. Neisseria gonorrhoeae Population Genomics: Use of the Gonococcal Core Genome to Improve Surveillance of Antimicrobial Resistance. J Infect Dis. 2020. doi: 10.1093/infdis/jiaa002. PubMed PMID: 32163580.

14. Harris SR, Cole MJ, Spiteri G, Sanchez-Buso L, Golparian D, Jacobsson S, et al. Public health surveillance of multidrug-resistant clones of Neisseria gonorrhoeae in Europe: a genomic survey. Lancet Infect Dis. 2018;18(7):758–68. doi: 10.1016/S1473-3099(18)30225-1 PubMed PMID: 29776807; PubMed Central PMCID: PMCPMC6010626.

15. Sanchez-Buso L, Golparian D, Corander J, Grad YH, Ohnishi M, Flemming R, et al. The impact of antimicrobials on gonococcal evolution. Nat Microbiol. 2019;4(11):1941–50. doi: 10.1038/s41564-019-0501-y. PubMed PMID: 31358980; PubMed Central PMCID: PMCPMC6817357.

16. Tanaka M, Nakayama H, Haraoka M, Saika T. Antimicrobial resistance of Neisseria gonorrhoeae and high prevalence of ciprofloxacin-resistant isolates in Japan, 1993 to 1998. J Clin Microbiol. 2000;38(2):521–5. PubMed PMID: 10655338; PubMed Central PMCID: PMCPMC86137.

17. Deguchi T, Yasuda M, Yokoi S, Ishida K, Ito M, Ishihara S, et al. Treatment of uncomplicated gonococcal urethritis by double-dosing of 200 mg cefixime at a 6-h interval. J Infect Chemother. 2003;9(1):35–9. doi: 10.1007/s10156-002-0204-8. PubMed PMID: 12673405.

18. Arnold BJ, Gutmann MU, Grad YH, Sheppard SK, Corander J, Lipsitch M, et al. Weak Epistasis May Drive Adaptation in Recombining Bacteria. Genetics. 2018;208(3):1247–60. doi: 10.1534/genetics.117.300662. PubMed PMID: 29330348; PubMed Central PMCID: PMCPMC5844334.

19. Grad YH, Kirkcaldy RD, Trees D, Dordel J, Harris SR, Goldstein E, et al. Genomic epidemiology of *Neisseria gonorrhoeae* with reduced susceptibility to cefixime in the USA: a retrospective observational study. Lancet Infect Dis. 2014;14(3):220–6. doi: 10.1016/S1473-3099(13)70693-5. PubMed PMID: 24462211; PubMed Central PMCID: PMCPMC4030102.

20. Golparian D, Harris SR, Sanchez-Buso L, Hoffmann S, Shafer WM, Bentley SD, et al. Genomic evolution of Neisseria gonorrhoeae since the preantibiotic era (1928-2013): antimicrobial use/misuse selects for resistance and drives evolution. BMC Genomics. 2020;21(1):116. doi: 10.1186/s12864-020-6511-6. PubMed PMID: 32013864; PubMed Central PMCID: PMCPMC6998845.

21. Costa-Lourenco A, Abrams AJ, Dos Santos KTB, Argentino ICV, Coelho-Souza T, Canine MCA, et al. Phylogeny and antimicrobial resistance in Neisseria gonorrhoeae isolates from Rio de Janeiro, Brazil. Infect Genet Evol. 2018;58:157–63. doi: 10.1016/j.meegid.2017.12.003. PubMed PMID: 29225148; PubMed Central PMCID: PMCPMC6932625.

22. Demczuk W, Lynch T, Martin I, Van Domselaar G, Graham M, Bharat A, et al. Whole-genome phylogenomic heterogeneity of Neisseria gonorrhoeae isolates with decreased cephalosporin susceptibility collected in Canada between 1989 and 2013. J Clin Microbiol. 2015;53(1):191–200. doi: 10.1128/JCM.02589-14. PubMed PMID: 25378573; PubMed Central PMCID: PMCPMC4290921.

23. Demczuk W, Martin I, Peterson S, Bharat A, Van Domselaar G, Graham M, et al. Genomic Epidemiology and Molecular Resistance Mechanisms of Azithromycin-Resistant Neisseria gonorrhoeae in Canada from 1997 to 2014. J Clin Microbiol. 2016;54(5):1304–13. doi: 10.1128/JCM.03195-15. PubMed PMID: 26935729; PubMed Central PMCID: PMCPMC4844716.

24. Eyre DW, De Silva D, Cole K, Peters J, Cole MJ, Grad YH, et al. WGS to predict antibiotic MICs for Neisseria gonorrhoeae. J Antimicrob Chemother. 2017;72(7):1937–47. doi: 10.1093/jac/dkx067. PubMed PMID: 28333355; PubMed Central PMCID: PMCPMC5890716.

25. Ezewudo MN, Joseph SJ, Castillo-Ramirez S, Dean D, Del Rio C, Didelot X, et al. Population structure of Neisseria gonorrhoeae based on whole genome data and its relationship with antibiotic resistance. PeerJ. 2015;3:e806. doi: 10.7717/peerj.806. PubMed PMID: 25780762; PubMed Central PMCID: PMCPMC4358642.

26. Kwong JC, Chow EPF, Stevens K, Stinear TP, Seemann T, Fairley CK, et al. Whole-genome sequencing reveals transmission of gonococcal antibiotic resistance among men who have sex with men: an observational study. Sex Transm Infect. 2018;94(2):151–7. doi: 10.1136/sextrans-2017-053287. PubMed PMID: 29247013; PubMed Central PMCID: PMCPMC5870456.

27. Lee RS, Seemann T, Heffernan H, Kwong JC, Goncalves da Silva A, Carter GP, et al. Genomic epidemiology and antimicrobial resistance of Neisseria gonorrhoeae in New Zealand. J Antimicrob Chemother. 2018;73(2):353–64. doi: 10.1093/jac/dkx405. PubMed PMID: 29182725; PubMed Central PMCID: PMCPMC5890773.

28. Ryan L, Golparian D, Fennelly N, Rose L, Walsh P, Lawlor B, et al. Antimicrobial resistance and molecular epidemiology using whole-genome sequencing of Neisseria gonorrhoeae in Ireland, 2014-2016: focus on extended-spectrum cephalosporins and azithromycin. Eur J Clin Microbiol Infect Dis. 2018;37(9):1661–72. doi: 10.1007/s10096-018-3296-5. PubMed PMID: 29882175.

29. Ma KC, Mortimer TD, Hicks AL, Wheeler NE, Sanchez-Buso L, Golparian D, et al. Adaptation to the cervical environment is associated with increased antibiotic susceptibility in Neisseria gonorrhoeae. Nat Commun. 2020;11(1):4126. doi: 10.1038/s41467-020-17980-1. PubMed PMID: 32807804.

30. Wiegand I, Hilpert K, Hancock RE. Agar and broth dilution methods to determine the minimal inhibitory concentration (MIC) of antimicrobial substances. Nat Protoc. 2008;3(2):163–75. doi: 10.1038/nprot.2007.521. PubMed PMID: 18274517.

31. Shigemura K, Okada H, Shirakawa T, Tanaka K, Arakawa S, Kinoshita S, et al. Susceptibilities of Neisseria gonorrhoeae to fluoroquinolones and other antimicrobial agents in Hyogo and Osaka, Japan. Sex Transm Infect. 2004;80(2):105–7. doi: 10.1136/sti.2003.006908. PubMed PMID: 15054169; PubMed Central PMCID: PMCPMC1744816.

32. Bankevich A, Nurk S, Antipov D, Gurevich AA, Dvorkin M, Kulikov AS, et al. SPAdes: a new genome assembly algorithm and its applications to single-cell sequencing. J Comput Biol. 2012;19(5):455–77. doi: 10.1089/cmb.2012.0021. PubMed PMID: 22506599; PubMed Central PMCID: PMCPMC3342519.

33. Seemann T. Shovill: faster SPAdes assembly of Illumina reads 2018 [cited 2020 June 21]. Available from: https://github.com/tseemann/shovill.

34. Koren S, Walenz BP, Berlin K, Miller JR, Bergman NH, Phillippy AM. Canu: scalable and accurate long-read assembly via adaptive k-mer weighting and repeat separation. Genome Res. 2017;27(5):722–36. doi: 10.1101/gr.215087.116. PubMed PMID: 28298431; PubMed Central PMCID: PMCPMC5411767.

35. Walker BJ, Abeel T, Shea T, Priest M, Abouelliel A, Sakthikumar S, et al. Pilon: an integrated tool for comprehensive microbial variant detection and genome assembly improvement. PLoS One. 2014;9(11):e112963. doi: 10.1371/journal.pone.0112963. PubMed PMID: 25409509; PubMed Central PMCID: PMCPMC4237348.

36. Langmead B, Salzberg SL. Fast gapped-read alignment with Bowtie 2. Nat Methods. 2012;9(4):357–9. doi: 10.1038/nmeth.1923. PubMed PMID: 22388286; PubMed Central PMCID: PMCPMC3322381.

37. Hunt M, Silva ND, Otto TD, Parkhill J, Keane JA, Harris SR. Circlator: automated circularization of genome assemblies using long sequencing reads. Genome Biol. 2015;16:294. doi: 10.1186/s13059-015-0849-0. PubMed PMID: 26714481; PubMed Central PMCID: PMCPMC4699355.

38. Seemann T. Prokka: rapid prokaryotic genome annotation. Bioinformatics. 2014;30(14):2068–9. doi: 10.1093/bioinformatics/btu153. PubMed PMID: 24642063.

39. Zhang G, Leclercq SO, Tian J, Wang C, Yahara K, Ai G, et al. A new subclass of intrinsic aminoglycoside nucleotidyltransferases, ANT(3”)-II, is horizontally transferred among Acinetobacter spp. by homologous recombination. PLoS Genet. 2017;13(2):e1006602. doi: 10.1371/journal.pgen.1006602. PubMed PMID: 28152054.

40. Darling AE, Mau B, Perna NT. progressiveMauve: multiple genome alignment with gene gain, loss and rearrangement. PloS one. 2010;5(6):e11147. doi: 10.1371/journal.pone.0011147. PubMed PMID: 20593022; PubMed Central PMCID: PMC2892488.

41. Unemo M, Golparian D, Sanchez-Buso L, Grad Y, Jacobsson S, Ohnishi M, et al. The novel 2016 WHO *Neisseria gonorrhoeae* reference strains for global quality assurance of laboratory investigations: phenotypic, genetic and reference genome characterization. J Antimicrob Chemother. 2016;71(11):3096–108. doi: 10.1093/jac/dkw288. PubMed PMID: 27432602; PubMed Central PMCID: PMCPMC5079299.

42. Didelot X, Pang B, Zhou Z, McCann A, Ni P, Li D, et al. The role of China in the global spread of the current cholera pandemic. PLoS Genet. 2015;11(3):e1005072. doi: 10.1371/journal.pgen.1005072. PubMed PMID: 25768799; PubMed Central PMCID: PMCPMC4358972.

43. Kurtz S, Phillippy A, Delcher AL, Smoot M, Shumway M, Antonescu C, et al. Versatile and open software for comparing large genomes. Genome Biol. 2004;5(2):R12. doi: 10.1186/gb-2004-5-2-r12. PubMed PMID: 14759262; PubMed Central PMCID: PMCPMC395750.

44. Guindon S, Dufayard JF, Lefort V, Anisimova M, Hordijk W, Gascuel O. New algorithms and methods to estimate maximum-likelihood phylogenies: assessing the performance of PhyML 3.0. Systematic biology. 2010;59(3):307–21. doi: 10.1093/sysbio/syq010. PubMed PMID: 20525638.

45. Stamatakis A. RAxML version 8: a tool for phylogenetic analysis and post-analysis of large phylogenies. Bioinformatics. 2014;30(9):1312–3. doi: 10.1093/bioinformatics/btu033. PubMed PMID: 24451623; PubMed Central PMCID: PMCPMC3998144.

46. Didelot X, Wilson DJ. ClonalFrameML: efficient inference of recombination in whole bacterial genomes. PLoS Comput Biol. 2015;11(2):e1004041. doi: 10.1371/journal.pcbi.1004041. PubMed PMID: 25675341; PubMed Central PMCID: PMCPMC4326465.

47. Rambaut A, Lam TT, Carvalho LM, Pybus OG. Exploring the temporal structure of heterochronous sequences using TempEst (formerly Path-O-Gen). Virus Evolution,. 2016;2(1):vew007.

48. Didelot X, Croucher NJ, Bentley SD, Harris SR, Wilson DJ. Bayesian inference of ancestral dates on bacterial phylogenetic trees. Nucleic Acids Res. 2018;46(22):e134. doi: 10.1093/nar/gky783. PubMed PMID: 30184106; PubMed Central PMCID: PMCPMC6294524.

49. Demczuk W, Sidhu S, Unemo M, Whiley DM, Allen VG, Dillon JR, et al. Neisseria gonorrhoeae Sequence Typing for Antimicrobial Resistance, a Novel Antimicrobial Resistance Multilocus Typing Scheme for Tracking Global Dissemination of N. gonorrhoeae Strains. J Clin Microbiol. 2017;55(5):1454–68. doi: 10.1128/JCM.00100-17. PubMed PMID: 28228492; PubMed Central PMCID: PMCPMC5405263.

50. Zankari E, Allesoe R, Joensen KG, Cavaco LM, Lund O, Aarestrup FM. PointFinder: a novel web tool for WGS-based detection of antimicrobial resistance associated with chromosomal point mutations in bacterial pathogens. J Antimicrob Chemother. 2017;72(10):2764–8. doi: 10.1093/jac/dkx217. PubMed PMID: 29091202; PubMed Central PMCID: PMCPMC5890747.

51. Hadfield J, Croucher NJ, Goater RJ, Abudahab K, Aanensen DM, Harris SR. Phandango: an interactive viewer for bacterial population genomics. bioRxiv. 2017.

52. Katoh K, Standley DM. MAFFT multiple sequence alignment software version 7: improvements in performance and usability. Mol Biol Evol. 2013;30(4):772–80. doi: 10.1093/molbev/mst010. PubMed PMID: 23329690; PubMed Central PMCID: PMCPMC3603318.

53. Waterhouse AM, Procter JB, Martin DM, Clamp M, Barton GJ. Jalview Version 2--a multiple sequence alignment editor and analysis workbench. Bioinformatics. 2009;25(9):1189–91. doi: 10.1093/bioinformatics/btp033. PubMed PMID: 19151095; PubMed Central PMCID: PMCPMC2672624.

54. Shimuta K, Unemo M, Nakayama S, Morita-Ishihara T, Dorin M, Kawahata T, et al. Antimicrobial resistance and molecular typing of *Neisseria gonorrhoeae* isolates in Kyoto and Osaka, Japan, 2010 to 2012: intensified surveillance after identification of the first strain (H041) with high-level ceftriaxone resistance. Antimicrob Agents Chemother. 2013;57(11):5225–32. doi: 10.1128/AAC.01295-13. PubMed PMID: 23939890; PubMed Central PMCID: PMCPMC3811299.

